# *Plasmodium* actin-like proteins 3 and 5a are essential for subsequent steps of mosquito infection

**DOI:** 10.64898/2026.05.22.727081

**Authors:** Caroline Busse, Yukino Kobayashi, Andrea Diers, Annika M. Binder, Friedrich Frischknecht, Ross G. Douglas

## Abstract

Actin superfamily members are critical for the biology of eukaryotes and archaea. Actin-related proteins (Arps) are a subgroup within the actin superfamily and play essential roles in trafficking, replication and motility. The genome of the malaria parasite *Plasmodium* contains a set of Arps unique to apicomplexans, termed actin-like proteins (Alps). However, the importance and specific roles of many of these Alps in *Plasmodium* progression are not yet understood. Here, we determined the functional contribution of *Plasmodium berghei* Alp3 and Alp5a (recently relabelled as Arp3) by generation of knock-out (KO) lines and their subsequent characterisation across different life cycle stages. Deletion of either Alp did not affect blood stage growth, gametogenesis and ookinete gliding motility. However, deletion of Alp5a lead to smaller and fewer oocysts as well as severely impaired sporozoite formation. The Alp3KO line had highly reduced oocyst loads compared to wild-type parasites. This striking decrease was due to impaired ookinete penetration of the mosquito midgut epithelium. Our study shows that both Alp3 and Alp5a are indispensable for *Plasmodium* transmission at different steps of initial mosquito infection, provides insights into the role of specific unique members of the actin superfamily during parasite progression and the requirements for efficient midgut penetration.

## Introduction

Malaria is a deadly tropical disease caused by parasites of the genus *Plasmodium* (World Health Organization 2025). During their life cycle, the parasites need to infect and adapt to very different host environments. This requires active motility as well as transcriptional, proteomic, metabolic and morphological changes in order to facilitate effective parasite replication and spread (Douglas et al. 2015; Cowman et al. 2016). After a blood meal, gametocytes transform into male and female gametes in the mosquito bolus that fuse to form a zygote which subsequently develops into an ookinete (Angrisano et al. 2012; Darif et al. 2025). Ookinetes actively move by gliding motility and penetrate the mosquito’s midgut epithelium where they transform into oocysts, which mature at the basal lamina and produce hundreds of sporozoites per oocyst (Baton and Ranford-Cartwright 2005; Douglas et al. 2015; Singer and Frischknecht 2023). These sporozoite egress into the haemocoel and then passively float through the haemolymph until invasion of the salivary glands for onward transmission (Douglas et al. 2015; Singer and Frischknecht 2023). Infection of the mosquito midgut represents one of the largest bottlenecks in the *Plasmodium* life cycle. Due to difficulties in culturing oocysts *in vitro*, and the relatively rapid nature of the infection process, little is known about this crucial mosquito stage and thus the proteins and protein dynamics required for these steps are still poorly understood (Smith and Barillas-Mury 2016; Hentzschel and Frischknecht 2022; Bailey et al. 2026).

The actin superfamily consists of multiple members, including conventional actin and actin-related proteins (Arps) (Kabsch and Holmes 1995; Hurley 1996). Arps are highly sequence divergent in comparison to conventional actin and are involved in processes such as intracellular trafficking, actin dynamics regulation and chromatin remodelling (Boyer and Peterson 2000; Oma and Harata 2011). Interestingly, *Plasmodium* has a reduced set of assigned classical Arp homologues (Gordon and Sibley 2005). The *Plasmodium* genome nonetheless contains a subgroup of six Arps found only in Apicomplexa, known as actin-like proteins (Alps), which are highly sequence divergent from the classical Arps (Gordon and Sibley 2005). Recently two Alps (Alp5a and Alp5b) were found to be part of an unusual Arp2/3 complex (Hentzschel et al. 2025). In general, very little is known about Alp function in apicomplexan biology. Here, we investigated the roles of two particular Alps, Alp3 and Alp5a (Arp3), in the biology of *Plasmodium*.

As members of the actin superfamily, Alp3 and Alp5a are predicted to share the conserved actin-fold core (Figure S1A). However, both Alps are significantly larger than conventional *Plasmodium* actin (539, and 574 amino acid residues for Alp3 and Alp5a respectively, compared to 376 residues for actin 1) (Figure S1A) and contain large insertions, many of which are located on the periphery. In terms of amino acid sequence, these Alps are highly divergent from *Plasmodium* actin 1 (13% identity and 28% similarity for Alp3 as well as 14% identity and 25% similarity for Alp5a) (Figure S1B, C). Through reverse genetics and imaging approaches, we identified key roles of these two Alps in ookinete penetration of the midgut epithelium and oocyst development.

## Results

### Alp3 and Alp5a are not required for asexual blood-stage growth, male gametogenesis and ookinete gliding motility

To investigate the roles of Alp3 and Alp5a in vertebrate and mosquito hosts, knock-out (KO) lines were generated in the rodent model *Plasmodium berghei*. Linearised knock-out vectors were transfected into schizonts (Figure S2). During limiting dilution, asexual blood-stage growth of these knock-out parasites was investigated. Relative growth rates were comparable to the complementation control, indicating that deletion of these Alps did not lead to strong impacts on blood-stage parasites (Figure 1A). These knock-out lines were then used to investigate the Alps’ role in the mosquito host.

**Figure 1:**
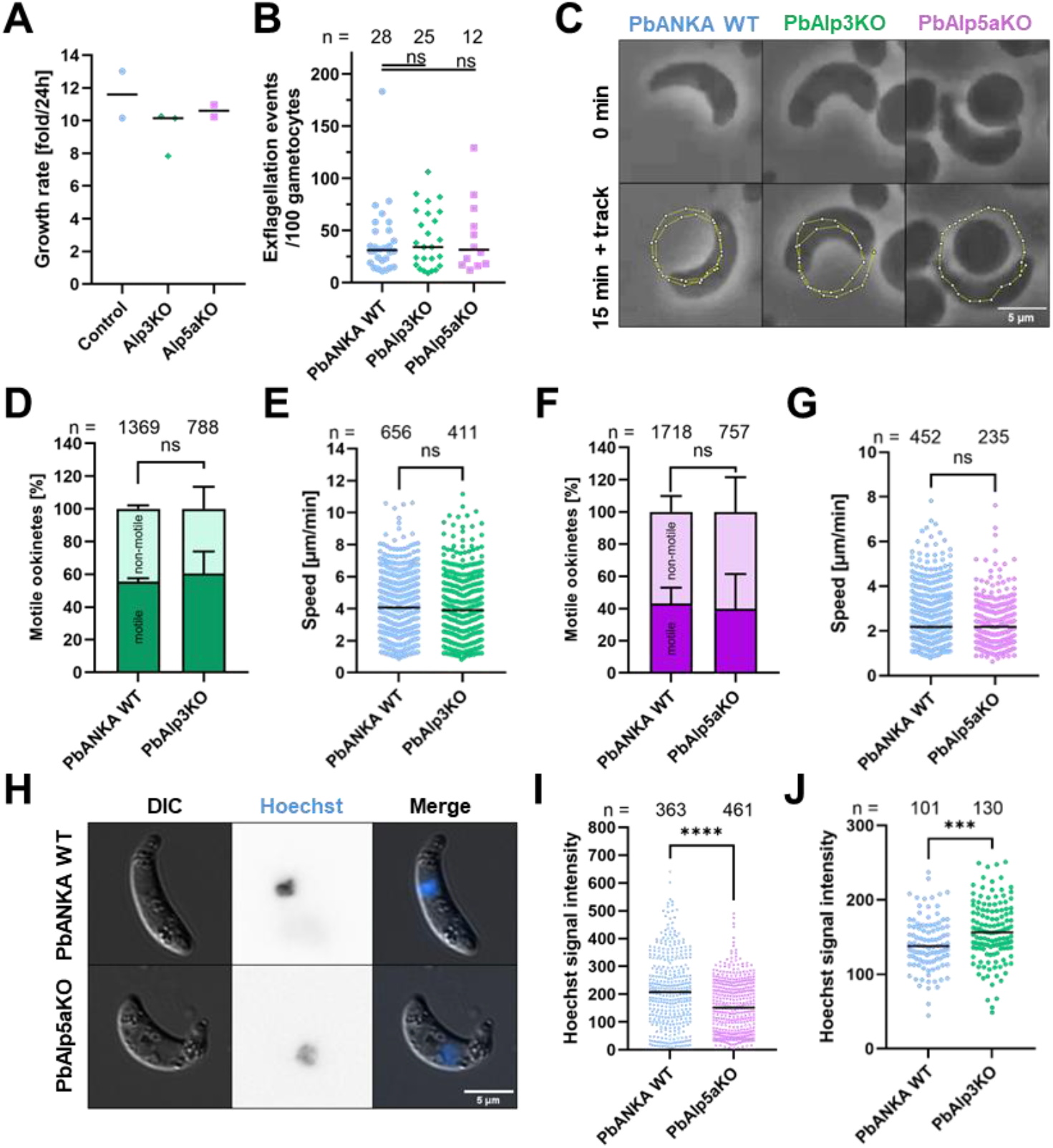
Alp3 and Alp5a are not required for blood stage growth, exflagellation or ookinete motility. **(A)** Knock-out (KO) of *P. berghei* Alp3 and Alp5a does not affect asexual blood-stage growth compared to control parasites. **(B)** Knock-out of Alp3 and Alp5a does not affect exflagellation of male gametocytes compared to PbANKA wild-type (WT). **(C)** Alp3KO and Alp5aKO ookinetes move in a circular pattern. Movement was tracked over 15 minutes. **(D, E)** Motile population and gliding speed of Alp3KO ookinetes is comparable to wild-type. **(F, G)** Motile population and gliding speed of Alp5aKO ookinetes is comparable to wild-type. **(H)** Hoechst signal is reduced in Alp5aKO ookinetes. **(I)** Alp5aKO ookinete nuclear Hoechst signal intensity is reduced by 25%. **(J)** Ookinete Hoechst signal intensity is not reduced in the Alp3KO line compared to wild-type, indicating no essential role at this stage of the life cycle. Statistics were performed using Fisher’s exact test (D, F) or Mann-Whitney test (E, G, I, J). ns = not significant; *** = p < 0.001; **** = p < 0.0001

To assess mosquito infection ability, we first investigated the Alps’ role in exflagellation of male gametocytes. Compared to wild-type parasites, no significant change in the number of exflagellation events was observed in Alp3KO and Alp5aKO parasites, highlighting the lack of Alp3 and Alp5a involvement in this process (Figure 1B). Ookinetes are highly motile parasite stages that employ active gliding motility to traverse the mosquito’s midgut epithelium to colonise a mosquito host. We thus explored the *in vitro* gliding motility of Alp3KO and Alp5aKO lines. Gliding motility of both knock-out ookinete lines were comparable to wild-type in terms of percentage of motile ookinetes and speeds (Figure 1C-G). Therefore, Alp3 and Alp5a do not play a major role in 2D *in vitro* ookinete gliding motility. Alp5a was recently identified as a member of an atypical Arp2/3 complex and knock-out of fellow complex members ARPC1 and Alp5b (Arp2) lead to a 33% reduction of DNA content in ookinetes compared to wild-type parasites (Hentzschel et al. 2025). To investigate the influence of Alp3 and Alp5a on the DNA content of ookinetes, Hoechst staining intensity was analysed. Nuclear staining intensity of Alp5aKO ookinetes was reduced by about 25% compared (Figures 1H, I), consistent with deletion of other complex members and a previous Alp5a knock-out (Hentzschel et al. 2025; Varshney et al. 2025). However, knock-out of Alp3 did not negatively affect ookinete Hoechst signal intensity (Figure 1J).

### Alp3 and Alp5a are essential for oocyst formation

Knock-out parasites were next investigated for their role in oocyst formation. Oocysts are sessile parasite midgut colonies which develop over the course of several days. Knock-out parasite lines were fed to *Anopheles stephensi* mosquitoes and infection rates analysed (Figure 2A). Alp3 and Alp5a knock-out lines were able to colonise mosquito midguts (Figure 2B) and the percentage of infected mosquitoes were comparable to the wild-type control (Figure 2C). However, Alp3KO-infected mosquitoes had approximately 99% reduced oocyst numbers (Figure 2D) and, in the case of the Alp5aKO line, total oocyst numbers were decreased by about 75% on day 11-12 after blood meal (Figure 2D). While the size of the few observed oocysts in the Alp3KO line was not negatively impacted, Alp5aKO resulted in reduced oocyst size by approximately 84% (Figure 2E). We generated complementation lines to investigate whether these reductions in oocyst numbers were caused by single-gene knock-outs (Figure S3). In both cases, oocyst numbers, and in case of the smaller oocysts for the Alp5aKO line, oocyst size was rescued in the complementation line (Figure 2D, E), indicating that Alp3 and Alp5a specifically are critical for oocyst formation.

**Figure 2:**
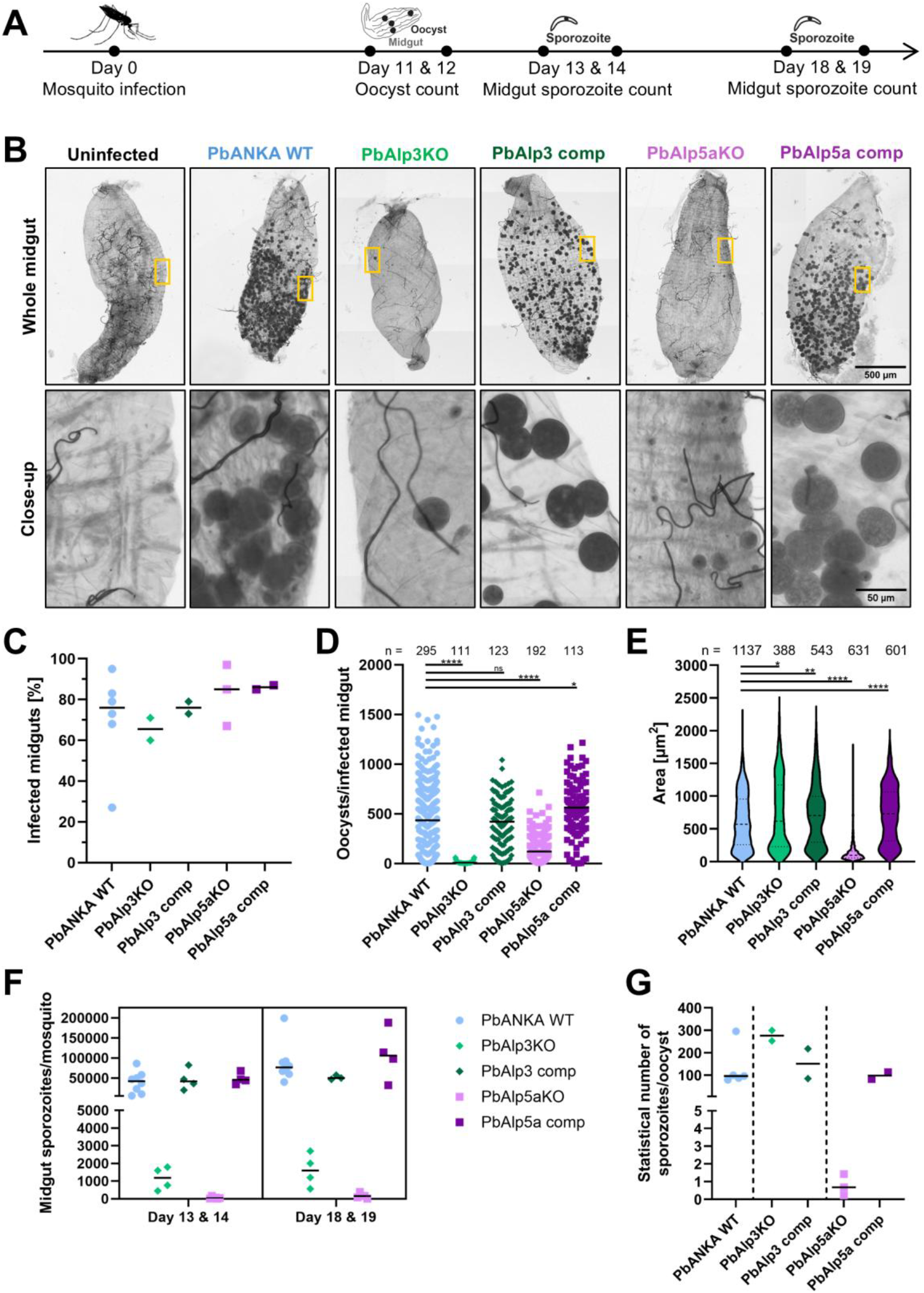
Alp3 and Alp5a are essential for oocyst formation. **(A)** Scheme of mosquito analysis timeline. **(B)** Microscopy images of *Anopheles* midguts stained with mercurochrome. **(C)** Knock-out of Alp3 and Alp5a did not affect infection rates of mosquito midguts. **(D)** Oocyst numbers in *Anopheles* midguts are reduced in the Alp3KO and Alp5aKO lines. **(E)** While knock-out of Alp3 had no effect on oocyst size, Alp5aKO oocysts were much smaller than wild-type. **(F)** Midgut sporozoite count is severely impaired in Alp3KO and Alp5aKO lines. **(G)** Normalisation of midgut sporozoite count (F) against oocysts per infected midgut (C, D) indicates an impairment of sporozoite formation in Alp5aKO oocysts. Statistics were performed using Mann-Whitney test. ns = not significant; * = p < 0.05; ** = p < 0.01; **** = p < 0.0001

### Alp5a, but not Alp3, is required for sporozoite formation

Hundreds of sporozoites develop per oocyst and egress into the haemocoel to reach and invade the salivary glands. Midgut sporozoites were quantified on days 13-14 and 18-19 post-infection. For the Alp3KO line, as expected due to very low total oocyst numbers, the quantity of Alp3KO midgut sporozoites was highly reduced compared to the wild-type control and was rescued by Alp3 complementation (Figure 2F). In case of the Alp5aKO line, total midgut sporozoite numbers were even lower with very few sporozoites observed (Figure 2F). This severe phenotype remained consistent over the course of five days and could be rescued by complementation with wild-type Alp5a. To determine whether the reduction in numbers was simply due to the reduction in total oocysts, sporozoite numbers were divided by the average number of oocysts. This ratio was comparable to wild-type in case of Alp3KO, indicating that the few oocysts present were able to sporulate efficiently (Figure 2G). However, in the case of the Alp5aKO line, the average number of sporozoites per oocyst was reduced to practically zero (Figure 2G). In summary, Alp3 is not required for sporulation and Alp5a is a critical factor for sporozoite development. Therefore, while both Alps are involved in oocyst formation, Alp5a is essential for oocyst maturation while Alp3 is required for events preceding oocyst formation.

### Alp3 is required for ookinete penetration of the midgut

In order for an oocyst to form, a *Plasmodium* ookinete needs to develop, penetrate the midgut epithelium, establish itself at the basal lamina and transform into an oocyst. To assess the functional role of Alp3 in these steps, we first investigated whether Alp3KO ookinetes can develop in the mosquito midgut. Fully mature Alp3KO ookinetes were readily observed in *in vitro* cultures (Figure 1C), and the blood bolus of infected mosquitoes dissected 24 hours after blood meal, indicating no major impact of Alp3 deletion on *in vivo* ookinete development (Figure 3A). Similar numbers of Alp3KO and wild-type ookinetes were found in the blood bolus (6000 and 6250 per microliter pelleted blood bolus for Alp3KO and wild-type, respectively). Next, we investigated whether Alp3KO ookinetes were able to penetrate the midgut epithelium. To this end, *Anopheles* midguts were dissected 24h post-infection, washed to remove any ookinetes remaining in the bolus and immuno-stained using a *Plasmodium*-specific α-HSP70 antibody. In the case of the Alp3KO, we detected a more than ten-fold reduction of parasites in the midgut epithelium (median of 92 and 7 parasites for wild-type and Alp3KO, respectively) (Figure 3B, C, Figure S4A), indicating a deficiency in epithelial layer penetration. This reduction was apparent regarding ookinetes and transforming ookinetes (tooks) as well as early oocysts (Figure 3D, E, Figure S4B). This reveals that Alp3 plays a critical role in ookinete penetration rather than in early oocyst development. Taken together, our data shows that two unique members of the actin superfamily, Alp3 and Alp5a, are critical for parasite progression in mosquitoes and are very likely involved in different processes.

**Figure 3:**
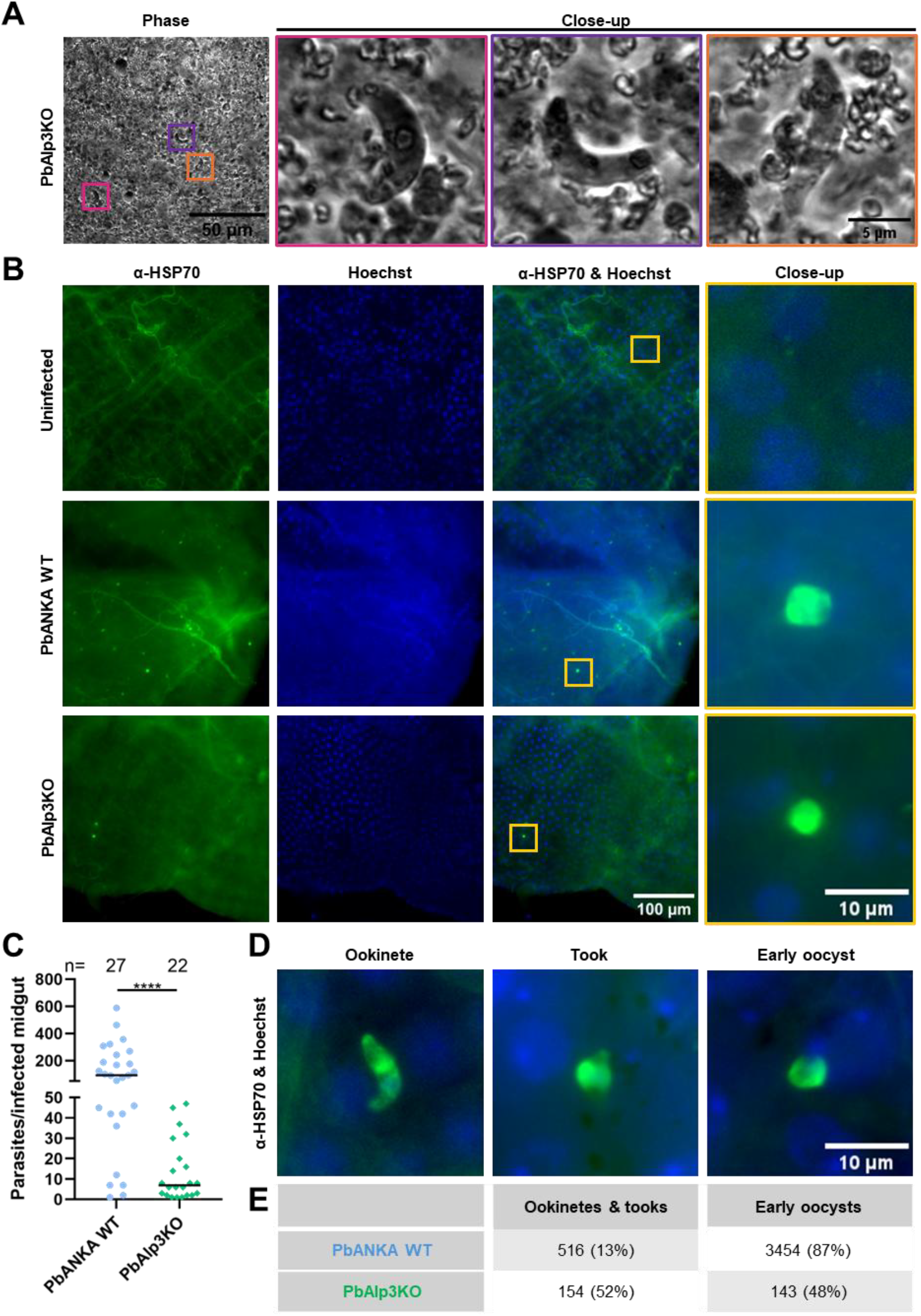
Alp3 is essential for midgut epithelium penetration. **(A)** PbAlp3KO ookinetes were present in blood bolus from mosquitoes dissected 24h post-infection. **(B)** Alp3KO parasites could be identified in the midgut epithelium by immuno-staining *Anopheles* midguts 24h post-infection using a *Plasmodium*-specific α-HSP70 antibody. **(C)** Quantification of parasites from (B) reveals that in the Alp3KO line, parasite numbers in the *Anopheles* midgut 24h post-infection are significantly reduced compared to wild-type. **(D)** Exemplary images of an ookinete and an early oocyst in the midgut epithelium. **(E)** Numbers of ookinetes and transitional stages (tooks) as well as early oocysts in the midgut were reduced in Alp3KO-infected midguts compared to wild-type. Statistics were performed using Mann-Whitney test. **** = p < 0.0001

## Discussion

Actin superfamily members are essential for multiple cellular processes across eukaryotes (Stoddard et al. 2017). *Plasmodium* conventional actin 1 is one of the most highly sequence divergent eukaryotic actins known, with this sequence divergence important for ookinete motility, smooth sporozoite motility and mosquito salivary gland penetration (Douglas et al. 2018; Yee et al. 2022; Kobayashi and Douglas 2025). The Arps subgroup has also been implicated or identified as essential for *Plasmodium* biology. Arp1 is required for efficient parasite blood stage growth (Bushell et al. 2017; Siden-Kiamos et al. 2010). Arp2 and 3 were originally thought to be absent in apicomplexans but, recently, Alp5b and Alp5a were discovered to be functioning in an atypical Arp2/3 complex that is critical for DNA segregation of activated male gametocytes (Hentzschel et al. 2025). Here, using reverse genetics and microscopy, we identified essential roles of Alp3 and Alp5a in *Plasmodium* transmission: Alp3 is essential for ookinete penetration of the mosquito midgut epithelium and Alp5a is necessary for oocyst maturation and sporozoite development.

Our knock-out study identified Alp5a as non-essential for blood stage growth, male gametogenesis and ookinete motility, yet as a critical factor for ookinete DNA content, oocyst development and subsequent sporozoite formation. Alp5a is expressed in a male gametocyte-specific manner (Howick et al. 2019) and, in a recent study identifying an atypical Arp2/3 complex in *Plasmodium*, knock-out of other complex members (ARPC1 and Alp5b) also lead to similar effects on DNA content and oocyst development. This further highlights the essential contributions of these Arp2/3 complex partner proteins in the function of the entire atypical complex (Hentzschel et al. 2025) and supports the suggestion that Alp5a is the functional homolog of Arp3. An Alp5a knock-out line was also recently published and a similar phenotype on oocyst number and size was observed (Varshney et al. 2025). Understanding the molecular interactions of this unique actin superfamily member will not only advance our knowledge of the structure-function relationships of Alp5a but also shed light on the molecular mechanisms of efficient DNA segregation in male gametocytes required for oocyst development. Considering the effect of Alp5aKO on DNA content in ookinetes, it is tempting to speculate that there could be a similar effect during sporulation in developing oocysts. Inducible knock-out methods or promoter swap approaches could be employed in the future to assess whether later life cycle steps require this complex for DNA segregation.

To our knowledge, this is the first in-depth investigation of the molecular role of Alp3, not only in *Plasmodium*, but in any apicomplexan. We found Alp3 to be essential for ookinete penetration of the mosquito midgut epithelium, a process that to date is still poorly understood (Hentzschel and Frischknecht 2022). Several proteins thus far have been shown to be involved in ookinete penetration of the midgut (Smith and Barillas-Mury 2016). Similar to the Alp3KO, knock-out of LIMP did not impair ookinete gliding motility but lead to severely reduced oocyst numbers (Egarter et al. 2021). Subtilisin-like protein PIMMS2 caused reduced midgut epithelium invasion of ookinetes (Ukegbu et al. 2017), CBP-O, a cyclic nucleotide binding protein in ookinetes, is also required for successful midgut penetration (Kwecka et al. 2025). Further, a quadruple mutant of the micronemal protein Akratin which, unlike the AkratinKO line, was able to exflagellate and form motile ookinetes but these ookinetes failed to penetrate the midgut epithelium (Kehrer et al. 2024). These findings highlight that the very short process of midgut epithelium penetration (Trisnadi and Barillas-Mury 2020) requires multiple protein contributors. In a recent study in which the transcriptome of early mosquito stages of the related *P. falciparum* was investigated, Alp3 was annotated as involved in cytoskeleton organisation. Furthermore, it was categorised in a gene cluster concerning rhoptry, nuclear migration and the inner membrane pellicle (Yan et al. 2025). Although we could not detect any defects in gliding motility in Alp3KO ookinetes, penetration of the mosquito midgut epithelium was severely impaired. Interestingly, an analogous observation was made in one of our previous studies on highly divergent *Plasmodium* actin residues in sporozoites, whereby mutant haemolymph sporozoites were generally able to glide to similar extents as wild-type controls, but were unable to efficiently penetrate salivary glands (Yee et al. 2022). The phenotype observed in the Alp3KO could be due to defects in the penetration machinery, possibly caused by slightly modified actin dynamics or reduced secretion of micronemal proteins, whereby sufficient dynamics or micronemal proteins are present to allow for gliding on a uniform glass surface but lack sufficient traction or force *in vivo* to penetrate the epithelial layer. Tagging strategies, especially when combined with microscopy techniques such as expansion microscopy, could reveal whether Alp3 is associated with the glideosome or micronemes and will shed light on the specific role Alp3 plays in this step of the parasite life cycle.

Mosquito infection is one of the largest bottlenecks in the *Plasmodium* life cycle. Our work has identified Alp3 as a key player in ookinete traversal of the midgut epithelium and Alp5a as essential for oocyst development (Figure 4). These findings highlight the involvement and requirement of unique actin superfamily members that are specialised in penetration of, and *Plasmodium* establishment in, the mosquito midgut.

**Figure 4:**
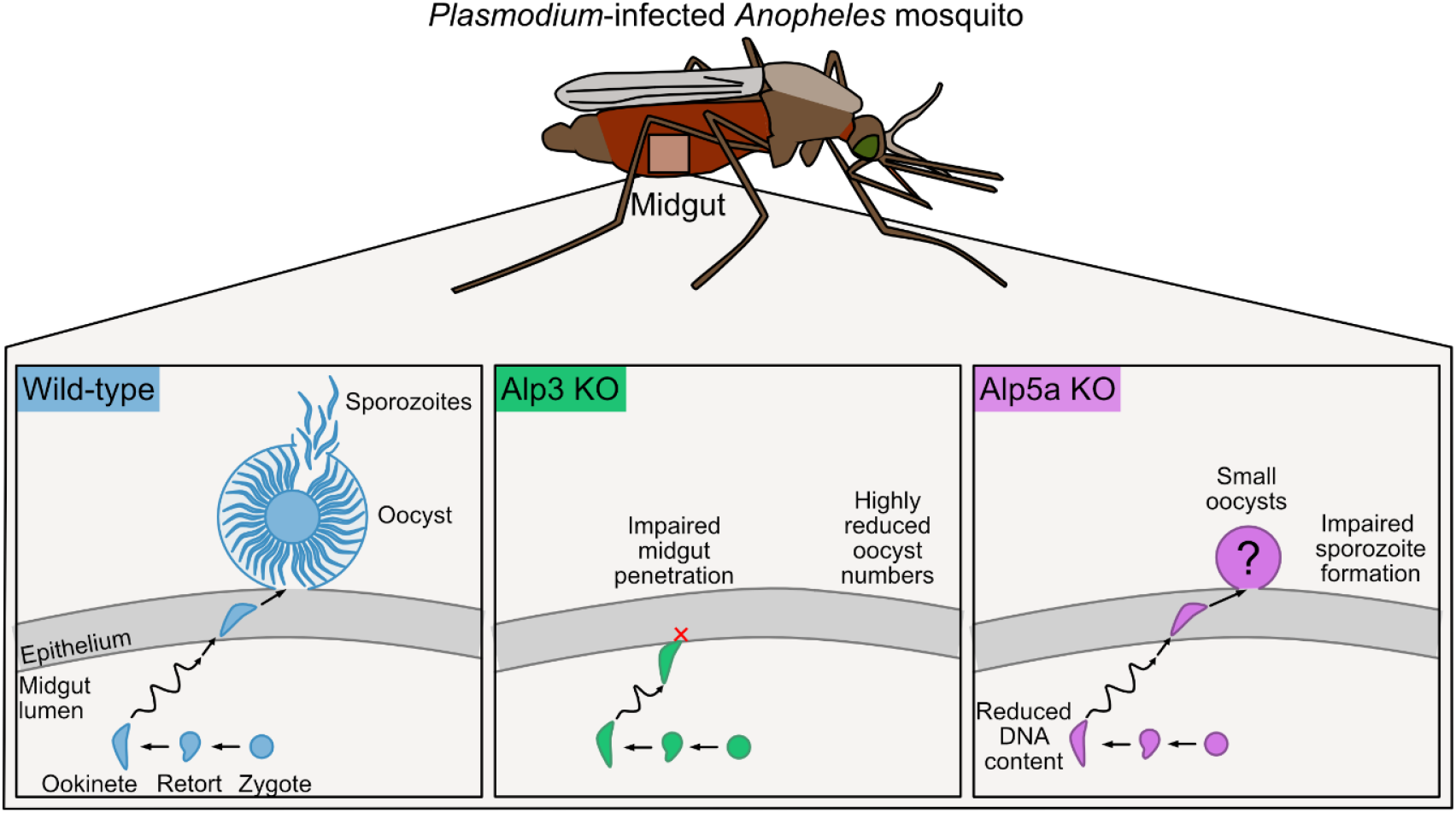
Graphical representation of the roles of Alp3 and Alp5a in *Plasmodium* transmission to mosquitoes.

## Experimental Procedures

### *In silico* structure predictions

*In-silico* structure predictions of *Plasmodium berghei* actin 1 (PBANKA_1459300), Alp3 (PBANKA_0212300) and Alp5a (PBANKA_0811800) were performed using AlphaFold2 (Bryant et al. 2022) and structures were coloured and modelled using open source PyMOL (Schroedinger). DNA and amino acid sequences were obtained from PlasmoDB (Bahl et al. 2003). Amino acid sequence identity and similarity were assessed using the EMBOSS Needle tool (EMBL, Madeira et al. 2024).

### Animal experiments and mosquito rearing

All animal work was approved by local authorities (Regierungspraesidium Giessen and Regierungspraesidium Karlsruhe) and performed according to the Federation of European Laboratory Animal Science Associations (FELASA) and Gesellschaft fuer Versuchstierkunde/Society for Laboratory Animal Science (GV-SOLAS) standard guidelines. Four-to six-week-old SWISS mice (Janvier Laboratories, France) were used for all animal experiments. *Anopheles stephensi* mosquitoes were reared and maintained according to standard breeding protocols (Benedict and Dotson 2015).

### Generation of knock-out and complementation parasite lines

Knock-out vectors of *Plasmodium berghei* Alp3 (PBANKA_0212300) and Alp5a (PBANKA_0811800) were kindly provided by the PlasmoGEM resource (Schwach et al. 2015). These vectors contain a *hDHFR* and *yfcu* genes, both under an eEF1 promotor. Transfection was performed as previously described (Janse et al. 2006) using a Nucleofector (Lonza) and parasites were selected using 0.07 mg/ml pyrimethamine that was added to the drinking water. Correct integration of DNA was investigated by harvesting mouse blood via cardiac puncture, gDNA purification using the DNeasy Blood & Tissue kit (Qiagen) and genotyping PCR (Table S1, Figure S2). Limiting dilution was performed to generate isogenic parasite lines. Asexual growth rates were calculated based on parasitaemia values from the limiting dilution as described before (Klug et al. 2016). To negatively select for parasites without selection cassette, 1.0-1.5 mg/ml 5-FC was added to the drinking water and limiting dilution was performed. Loss of selection cassette was confirmed via genotyping PCR (Table S1, Figure S3).

For complementation lines, wild-type Alp genes were obtained from *P. berghei* gDNA (Table S1) and cloned into Pb238 transfection vectors (Douglas et al. 2018) using BssHII and BamHI (NEB). These vectors were linearised using SacII and PmeI (NEB) and transfected as above into negatively selected KO parasites. Limiting dilution was performed and correct integration of the gene was confirmed via genotyping PCR (Table S1).

### Exflagellation assay

Swiss mice were infected with cryo-preserved *Plasmodium* parasites via intra-peritoneal injection. When parasitaemia reached 1.5-2%, blood was harvested via cardiac puncture and blood corresponding to 20 million parasites were transferred into a recipient mouse. Three days later, a drop of tail blood from an infected mouse was placed on a microscopy slide and covered with a coverslip. The slides were incubated at RT in the dark for 9 minutes. Exflagellation centres were quantified across multiple fields of view under a Zeiss AxioLab A1 light microscope using a 40x objective and phase contrast ring between minutes 9-14 post-activation. Red blood cells were counted in one field. Gametocytaemia was assessed by preparing a Giemsa-stained blood smear. Exflagellation centres were normalised against gametocyte numbers.

### Ookinete culture

Ookinete culture analysis was performed as described previously (Douglas et al. 2018; Yee et al. 2022). Briefly, mice were infected and fresh blood transfer was performed as described above. Three days later, if one or more exflagellation centres per microscope field could be observed, blood was collected and incubated in ookinete medium (250 ml RPMI1640 + HEPES + glutamine, 12.5 mg hypoxanthine, 2.5 ml penicillin/streptomycin (100x), 0.5 g sodium bicarbonate, 5.12 mg xanthurenic acid (100 μM), 16% (v/v) foetal bovine serum, pH 7.8) at 19°C for 21h in the dark. Development of ookinetes was confirmed by Giemsa smear of the culture. Ookinetes were enriched by adding a cushion of 10 ml 63% Nycodenz to 10 ml ookinete culture and centrifuging at 1000 rpm/25 min/RT without break. The ookinete-containing ring at the interface was collected and pelleted at 1000 rpm/10 min/RT with break. The pellet was resuspended in 1 ml ookinete medium.

For motility and speed analysis, 100-200 µl ookinete culture was centrifuged at 13000 rpm/10 sec/RT and 2 µl of the pellet with added on a microscopy slide, covered with a coverslip and sealed with paraffin wax. Ookinetes were imaged under an Axioskop 2 MOT (Zeiss) every 30 sec for 15 min using a 40x objective. Movies were analysed using FIJI (Schindelin et al. 2012), employing the manual tracking plug-in for speed analysis.

For ookinete DNA content analysis, PbAlp3KO and respective wild-type control ookinetes were fixed in 4% PFA/PBS for 10 min at RT in the dark and washed. In case of PbAlp5aKO and the respective wild-type control, ookinetes were not fixed. In both cases, ookinetes were stained with 10 µg/ml Hoechst 33342 (Thermo Fisher) in ookinete medium for 10 min at RT in the dark and centrifuged at 13000 rpm/10 sec/RT. The pellet was resuspended in ookinete medium, centrifuged again and 2 µl of the pellet were added to a microscopy slide, covered with a coverslip and sealed with paraffin wax. Ookinetes were imaged under an Axio Observer Z1 (Zeiss) using a 63x objective (DIC channel at 150 ms exposure and Hoechst 33258 channel at 25 ms exposure). Hoechst signal intensity was analysed using a FIJI macro as described previously (Hentzschel et al. 2025).

### Mosquito infection

For infections, a donor mouse was infected with cryo-preserved parasites and, upon reaching a parasitaemia of 1.5-2%, blood was harvested. Blood containing twenty million parasites were transferred into two recipient mice. When at least one exflagellation event per field could be detected, the mice were anaesthetised (100 mg/kg ketamine, 3 mg/kg xylazine), placed on a mosquito cage that had been starved for 3-6h and mosquitoes allowed to feed for 30 minutes. Mice were sacrificed via cervical dislocation directly after feeding. The mosquito cage was placed in an incubator at 21°C and 70% humidity and mosquitoes were maintained with 10% saccharose with 0.05% PABA and 0.1% NaCl solution.

### Characterisation of mosquito stage parasites

Mosquitoes were collected and placed on ice before being briefly dipped into 70% EtOH and transferred to a dish containing PBS. Prepared mosquitoes were then placed on a microscopy slide and midguts dissected under a SMZ1B stereo microscope (Nikon) in PBS. Midguts were collected in PBS on ice and further analysed.

For mercurochrome staining, midguts were dissected on day 11 and 12 post-infection and permeabilised with NP-40 (AppliChem, final concentration 1% in PBS) for 20 min at RT. NP40 solution was removed and midguts were stained in 1 ml 0.1% mercurochrome/PBS for 2h at RT. The samples were washed 4x with PBS before being aligned on a microscopy slide and covered with a coverslip. Stained midguts were imaged under a Axio Observer Z1 (Zeiss) using a 10x objective with a green filter. Infection rates, oocyst number and oocyst size were analysed using FIJI (Schindelin et al. 2012).

For midgut sporozoite counts, midguts were dissected into 50 µl PBS on day 13, 14, 18 and 19 post-infection. Dissected midguts were crushed with a pestle for 1 min and diluted with PBS before adding 10 µl of the sample onto a Neubauer counting chamber. Parasites were counted on a light microscope with phase contrast ring using a 40x objective.

For *in vivo* ookinete and early oocyst analysis, midguts of female *Anopheles* mosquitoes were dissected 24h post-infection and carefully sliced open longitudinally to remove the blood bolus. To analyse ookinete formation in the blood bolus, blood boluses of 15 infected midguts were collected and fixed with 4% PFA/PBS for 1h at RT and washed 4x in PBS. For detection of ookinetes, the samples were pelleted (7000 rpm/2 min/RT) and supernatant removed. 2 µl of the wet pellet was placed on a microscopy slide, covered with a coverslip and sealed with paraffin wax before imaging on a DM6 B microscope (Leica) using a 40x objective. Images were analysed using FIJI (Schindelin et al. 2012). To quantify ookinetes in the blood bolus, fixed blood bolus samples were pelleted by centrifugation and the supernatant was removed. 1 µl of the pellet was collected and added to 99 µl PBS. After thorough mixing, 10 µl of the sample were pipetted onto a Neubauer counting chamber. The number of ookinetes was assessed under an BX50 microscope (Olympus) with DIC filter using a 40x objective. For ookinete analysis in the midgut epithelium, empty midguts were fixed with 4% PFA/PBS for 1h at RT. Fixed midguts were washed 4x with PBS and permeabilised with 0.1% Triton-X-100/PBS for 10 min at RT. The samples were washed, blocked with 5% BSA/PBS for 1h at RT and stained overnight with a *Plasmodium*-specific α-HSP70 antibody. After another wash step, parasites were visualised using an α-rabbit antibody coupled with AlexaFluor 488 (Invitrogen) for 1h. After 30 min, Hoechst 33342 was added (Thermo Fisher, final concentration 1 µg/µl). The samples were washed again, and the midguts were gently placed on a microscopy slide, covered with a coverslip and sealed around the edges using paraffin wax. Imaging was performed using a DM6 B fluorescence microscope (Leica) using a 40x objective and an Axio Observer Z1 with an Apotome (Zeiss) using a 63x objective. Image analysis was performed using FIJI (Schindelin et al. 2012).

## Acknowledgements

We thank Franziska Hentzschel for helpful discussions and for providing the macro used for DNA content analysis. We also thank Jude Przyborski for kindly providing the *Plasmodium* HSP70 antibody. This work is supported by the German Research Foundation (DFG grant DO 2329/3-1).

## Publication bibliography

Angrisano, Fiona; Tan, Yan-Hong; Sturm, Angelika; McFadden, Geoffrey I.; Baum, Jake (2012): Malaria parasite colonisation of the mosquito midgut--placing the Plasmodium ookinete centre stage. In International Journal for Parasitology 42 (6), pp. 519–527. DOI: 10.1016/j.ijpara.2012.02.004.

Bahl, Amit; Brunk, Brian; Crabtree, Jonathan; Fraunholz, Martin J.; Gajria, Bindu; Grant, Gregory R. et al. (2003): PlasmoDB: the Plasmodium genome resource. A database integrating experimental and computational data. In Nucleic Acids Research 31 (1).

Bailey, Alexander J.; Vlachou, Dina; Christophides, George K. (2026): Oocyst: knowns and unknowns about the lengthiest life stage of the malaria parasite. In Open biology 16 (4). DOI: 10.1098/rsob.260009.

Baton, Luke A.; Ranford-Cartwright, Lisa C. (2005): How do malaria ookinetes cross the mosquito midgut wall? In Trends in parasitology 21 (1).

Benedict, Mark Q.; Dotson, Ellen M. (2015): Methods in Anopheles Research.

Boyer, Laurie A.; Peterson, Craig L. (2000): Actin-related proteins (Arps): conformational switches for chromatin-remodeling machines? In BioEssays 22 (7), pp. 666–672.

Bryant, Patrick; Pozzati, Gabriele; Elofsson, Arne (2022): Improved prediction of protein-protein interactions using AlphaFold2. In Nature communications 13 (1265).

Bushell, Ellen; Gomes, Ana Rita; Sanderson, Theo; Anar, Burcu; Girling, Gareth; Herd, Colin et al. (2017): Functional Profiling of a Plasmodium Genome Reveals an Abundance of Essential Genes. In Cell 170 (2), 260–272.e8. DOI: 10.1016/j.cell.2017.06.030.

Cowman, Alan F.; Healer, Julie; Marapana, Danushka; Marsh, Kevin (2016): Malaria: Biology and Disease. In Cell 167 (3), pp. 610–624. DOI: 10.1016/j.cell.2016.07.055.

Darif, Nedal; Rheinnecker, Marco; Hildenbrand, Kolja; Chookajorn, Thanat; Dorner, Lilian P.; Hériché, Jean-Karim et al. (2025): Cellular Hallmarks From Volume Electron Microscopy Reveal Developmental Progression of Plasmodium Ookinetes. In Advanced science (Weinheim, Baden-Wurttemberg, Germany), e08250. DOI: 10.1002/advs.202508250.

Douglas, Ross G.; Amino, Rogerio; Sinnis, Photini; Frischknecht, Freddy (2015): Active migration and passive transport of malaria parasites. In Trends in parasitology 31 (8), pp. 357–362. DOI: 10.1016/j.pt.2015.04.010.

Douglas, Ross G.; Nandekar, Prajwal; Aktories, Julia-Elisabeth; Kumar, Hirdesh; Weber, Rebekka; Sattler, Julia M. et al. (2018): Inter-subunit interactions drive divergent dynamics in mammalian and Plasmodium actin filaments. In PLoS biology 16 (7).

Egarter, Saskia; Santos, Jorge M.; Kehrer, Jessica; Sattler, Julia; Frischknecht, Friedrich; Mair, Gunnar R. (2021): Gliding motility protein LIMP promotes optimal mosquito midgut traversal and infection by Plasmodium berghei. In Molecular and biochemical parasitology 241.

Gordon, Jennifer L.; Sibley, L. David (2005): Comparative genome analysis reveals a conserved family of actin-like proteins in apicomplexan parasites. In BMC Genomics 6 (179).

Hentzschel, Franziska; Frischknecht, Friedrich (2022): Still enigmatic: Plasmodium oocysts 125 years after their discovery. In Trends in parasitology 38 (8), pp. 610–613. DOI: 10.1016/j.pt.2022.05.013.

Hentzschel, Franziska; Jewanski, David; Sokolowski, Yvonne; Agarwal, Pratika; Kraeft, Anna; Hildenbrand, Kolja et al. (2025): An atypical Arp2/3 complex is required for Plasmodium DNA segregation and malaria transmission. In Nat Microbiol 10 (7), pp. 1775–1790. DOI: 10.1038/s41564-025-02023-6.

Howick, Virginia M.; Russell, Andrew J. C.; Andrews, Tallulah; Heaton, Haynes; Reid, Adam J.; Natarajan, Kedar et al. (2019): The Malaria Cell Atlas: Single parasite transcriptomes across the complete Plasmodium life cycle. In Science (New York, N.Y.) 365 (6455). DOI: 10.1126/science.aaw2619.

Hurley, James H. (1996): The Sugar Kinase/Heat Shock Protein 70/Actin Superfamily: Implications of Conserved Structure for Mechanism. In Annu. Rev. Biophys. Biomol. 25, pp. 137–162.

Janse, Chris J.; Ramesar, Jai; Waters, Andrew P. (2006): High-efficiency transfection and drug selection of genetically transformed blood stages of the rodent malaria parasite Plasmodium berghei. In Nature Protocols 1 (1).

Kabsch, W.; Holmes, K. C. (1995): The actin fold. In FASEB journal: official publication of the Federation of American Societies for Experimental Biology 9 (2), pp. 167–174. DOI: 10.1096/fasebj.9.2.7781919.

Kehrer, Jessica; Pietsch, Emma; Ricken, Dominik; Strauss, Léanne; Heinze, Julia M.; Gilberger, Tim; Frischknecht, Friedrich (2024): APEX-based proximity labeling in Plasmodium identifies a membrane protein with dual functions during mosquito infection. In PLoS pathogens 20 (12), e1012788. DOI: 10.1371/journal.ppat.1012788.

Klug, Dennis; Mair, Gunnar R.; Frischknecht, Friedrich; Douglas, Ross G. (2016): A small mitochondrial protein present in myzozoans is essential for malaria transmission. In Open biology 6 (4), p. 160034. DOI: 10.1098/rsob.160034.

Kobayashi, Yukino; Douglas, Ross G. (2025): Highly divergent apicomplexan cytoskeletons provide additional models for actin biology. In The FEBS journal. DOI: 10.1111/febs.70263.

Kwecka, Dominika; Wang, Zhishuo; Eigminas, Edvardas; Liu, Lijia; Regan, Jennifer C.; Kim, Choel; Philip, Nisha (2025): A divergent cyclic nucleotide binding protein promotes Plasmodium ookinete infection of the mosquito. In PLoS pathogens 21 (9), e1013467. DOI: 10.1371/journal.ppat.1013467.

Madeira, Fábio,; Madhusoodanan, Nandana; Lee, Joonheung; Eusebi, Alberto; Niewielska, Ania; Tivey, Adrian R. N. et al. (2024): The EMBL-EBI Job Dispatcher sequence analysis tools framework in 2024. In Nucleic Acids Research 52 (W1), W521–W525. DOI: 10.1093/nar/gkae241.

Oma, Yukako; Harata, Masahiko (2011): Actin-related proteins localized in the nucleus. From discovery to novel roles in nuclear organization. In Nucleus 2 (1), pp. 38–46.

Schindelin, Johannes; Arganda-Carreras, Ignacio; Frise, Erwin; Kaynig, Verena; Longair, Mark; Pietzsch, Tobias et al. (2012): Fiji: an open-source platform for biological-image analysis. In Nature methods 9 (7), pp. 676–682. DOI: 10.1038/nmeth.2019.

Schwach, Frank; Bushell, Ellen; Gomes, Ana Rita; Anar, Burcu; Girling, Gareth; Herd, Colin et al. (2015): PlasmoGEM, a database supporting a community resource for large-scale experimental genetics in malaria parasites. In Nucleic Acids Research 43 (Database issue), D1176–82. DOI: 10.1093/nar/gku1143.

Siden-Kiamos, Inga; Schüler, Herwig; Liakopoulos, Dimitris; Louisa, Christos (2010): Arp1, an actin-related protein, in Plasmodium berghei. In Molecular and biochemical parasitology 173, pp. 88–96.

Singer, Mirko; Frischknecht, Friedrich (2023): Still running fast: Plasmodium ookinetes and sporozoites 125 years after their discovery. In Trends in parasitology 39 (12), pp. 991–995. DOI: 10.1016/j.pt.2023.09.015.

Smith, Ryan C.; Barillas-Mury, Carolina (2016): Plasmodium Oocysts: Overlooked Targets of Mosquito Immunity. In Trends in parasitology 32 (12), pp. 979–990. DOI: 10.1016/j.pt.2016.08.012.

Stoddard, Patrick R.; Williams, Tom A.; Garner, Ethan; Baum, Buzz (2017): Evolution of polymer formation within the actin superfamily. In Molecular Biology of the Cell 28.

Trisnadi, Nathanie; Barillas-Mury, Carolina (2020): Live In Vivo Imaging of Plasmodium Invasion of the Mosquito Midgut. In mSphere 5 (5).

Ukegbu, Chiamaka V.; Akinosoglou, Karolina A.; Christophides, George K.; Vlachou, Dina (2017): Plasmodium berghei PIMMS2 Promotes Ookinete Invasion of the Anopheles gambiae Mosquito Midgut. In Infection and immunity 85 (8). DOI: 10.1128/IAI.00139-17.

Varshney, Aastha; Pandey, Eisha Nirdosh; Mishra, Satish (2025): Plasmodium actin-like proteins are essential for DNA segregation during male gametogenesis and malaria transmission. In PLoS pathogens 21 (11), e1013687. DOI: 10.1371/journal.ppat.1013687.

World Health Organization (2025): World malaria report 2025: Addressing the threat of antimalarial drug resistance.

Yan, Yan; Verzier, Lisa H.; Cheung, Elaine; Appetecchia, Federico; March, Sandra; Craven, Ailsa R. et al. (2025): Mapping Plasmodium transitions and interactions in the Anopheles female. In Nature 648 (8093), pp. 451–458. DOI: 10.1038/s41586-025-09653-0.

Yee, Michelle; Walther, Tobias; Frischknecht, Friedrich; Douglas, Ross G. (2022): Divergent Plasmodium actin residues are essential for filament localization, mosquito salivary gland invasion and malaria transmission. In PLoS pathogens 18 (8).

