## Supplementary figures and tables for "*Plasmodium* actin-like proteins 3 and 5a are essential for subsequent steps of mosquito infection"

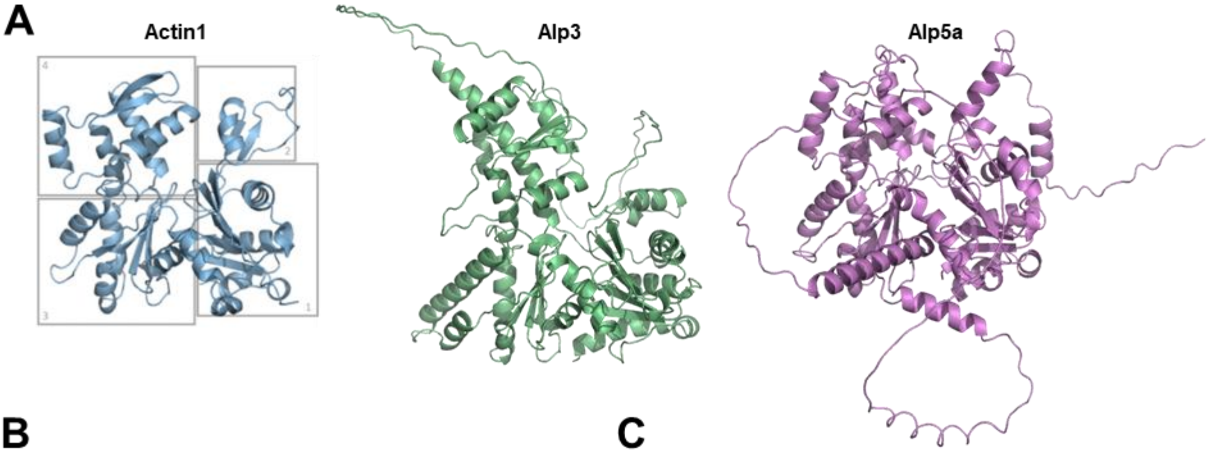


**B**

| Identity | Actin1 | Alp3 | Alp5a |
|---|---|---|---|
| Actin1 | 100 | 13 | 14 |
| Alp3 | 13 | 100 | 18 |
| Alp5a | 14 | 18 | 100 |

**C**

| Similarity | Actin1 | Alp3 | Alp5a |
|---|---|---|---|
| Actin1 | 100 | 28 | 25 |
| Alp3 | 28 | 100 | 30 |
| Alp5a | 25 | 30 | 100 |

**Figure S1: Alp3 and Alp5a are highly sequence divergent from actin and are predicted to have an actin fold. (A)** AlphaFold2 structure prediction of *P. berghei* actin 1, Alp3 and Alp5a. Gray outlines represent four subdomains which, in addition to a nucleotide binding cleft in the centre, are characteristic for the actin-fold. Alp3 and Alp5a share the conserved actin-fold core but have large peripheral insertions. **(B), (C)** Amino acid sequence identity [%] **(B)** and similarity [%] **(C)** are low between actin 1 and Alp3 or Alp5a from *P. berghei.*

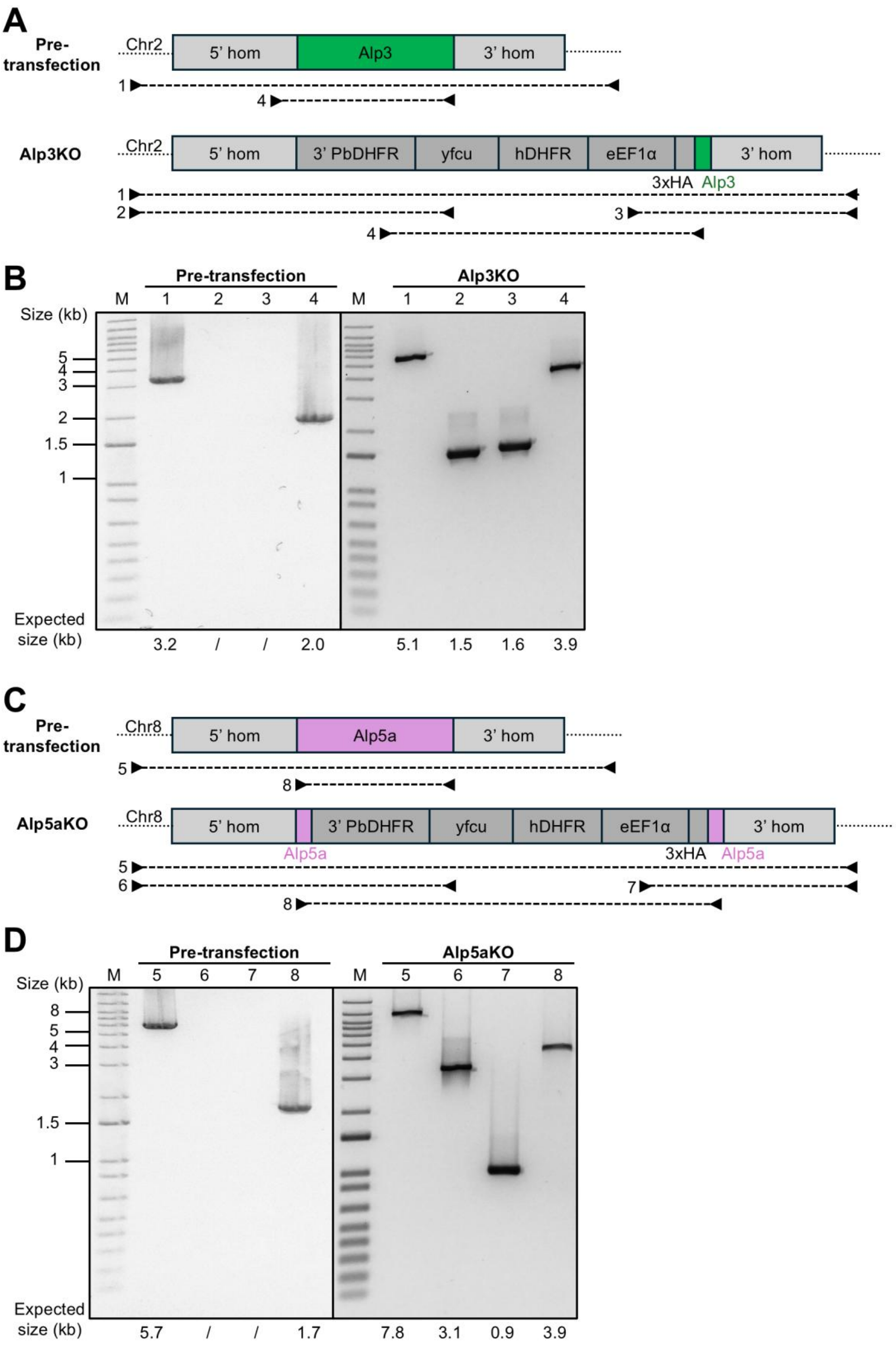


**Figure S2: Genotyping of *P. berghei* Alp3 and Alp5a knock-out lines. (A)** Genotyping strategy of Alp3 knock-out line. **(B)** Genotyping gel shows correct integration of the knock-out vector. **(C)** Genotyping strategy of Alp5a knock-out line.

**(D)** Genotyping gel shows correct integration of the knock-out vector. Chr = chromosome. eEF1α = eucaryotic translation elongation factor 1α. DHFR = dihydrofolate reductase. HA = haemagglutinin. Hom = homology region. Kb = kilo base pairs. KO = knock-out. ORF = open reading frame. WL = whole locus. Yfcu = yeast cytosine deaminase and uridyl-phosphoribosyltransferase.

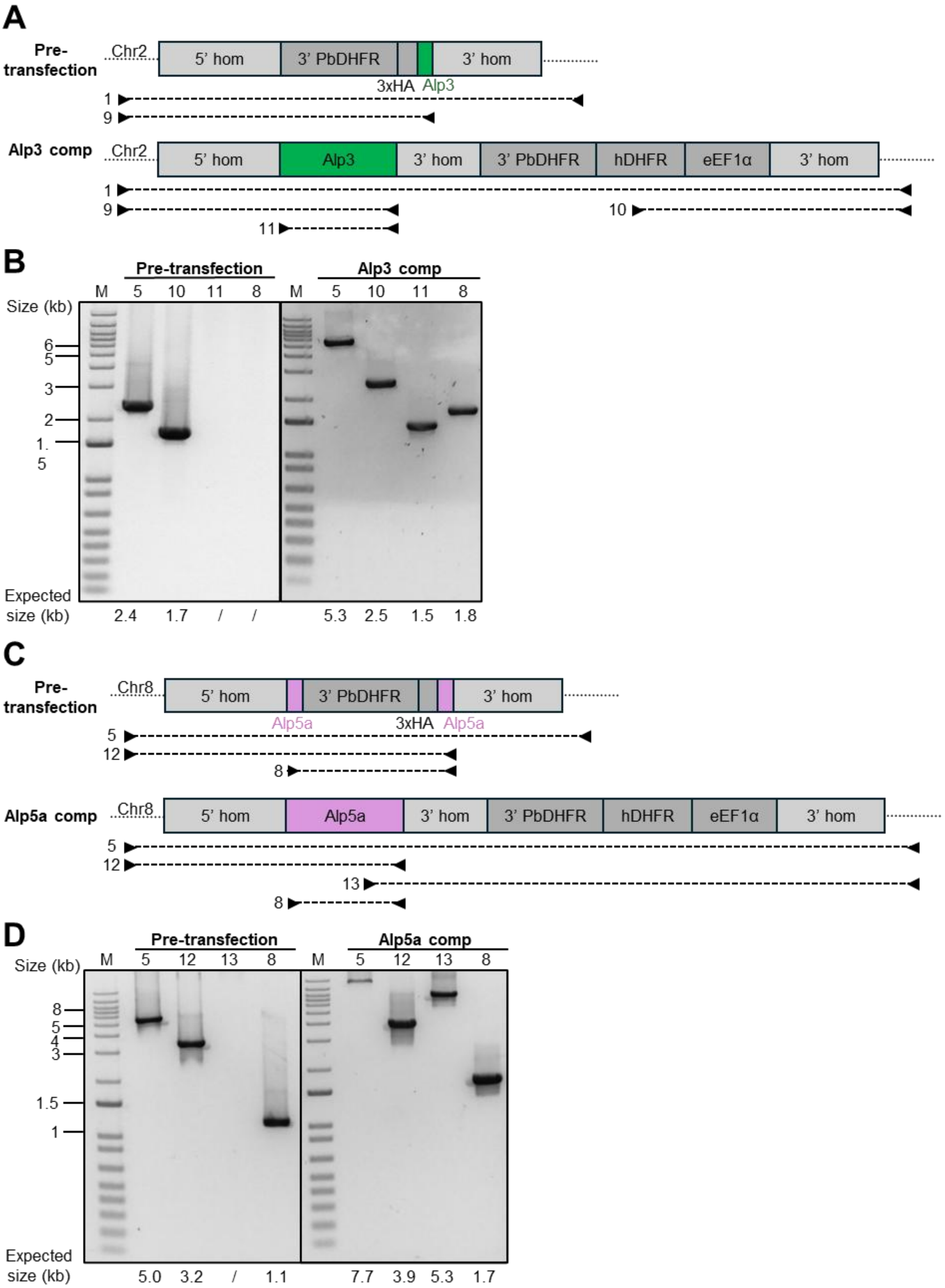


**Figure S3**: **Genotyping of *P. berghei* Alp3 and Alp5a complementation lines. (A)** Genotyping strategy of Alp3 complementation line. **(B)** Genotyping gel shows correct integration of the complementation vector. **(C)** Genotyping strategy of Alp5a complementation line. **(D)** Genotyping gel shows correct integration of the complementation vector. Chr = chromosome. Comp. = complementation. eEF1α = eucaryotic translation elongation factor 1α. DHFR = dihydrofolate reductase. HA = haemagglutinin. Hom = homology region. Kb = kilo base pairs. KO = knock-out. Neg. sel. = negatively selected. ORF = open reading frame. WL = whole locus. Yfcu = yeast cytosine deaminase and uridyl-phosphoribosyltransferase.

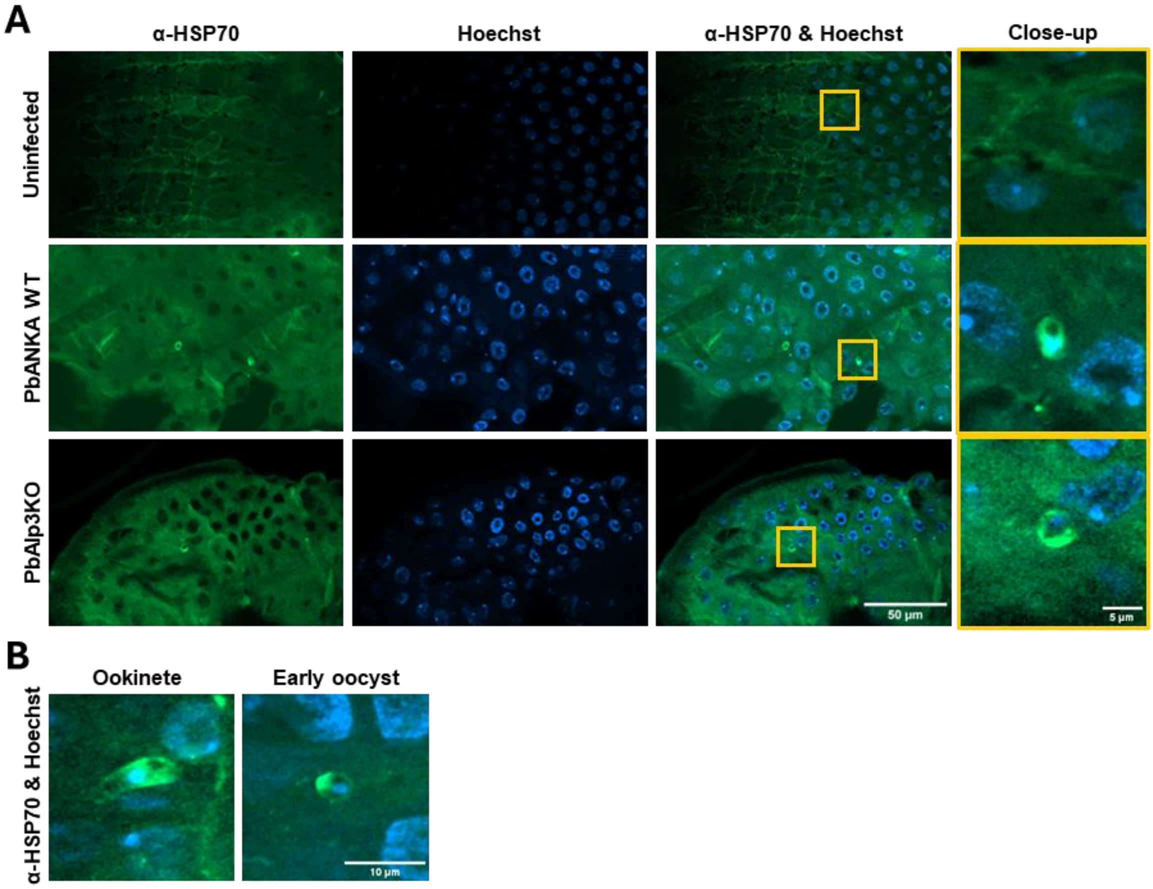


**Figure S4: Additional examples of fixed and stained *Plasmodium*-infected Anopheles midguts. (A)** Both wild-type and Alp3KO parasites could be detected in the midgut epithelium. Parasites were immuno-stained using an anti-HSP70 antibody and imaged with a fluorescence microscope using an Apotome (Zeiss). **(B)** Representative images of an ookinete and an early oocyst in the midgut epithelium.

**Table S1: Primers used for cloning and genotyping PCRs.**

| Gene | Direction | Primer sequence (5' - 3') |
|---|---|---|
| Alp3 | Forward | ATCACTTTTTCTTTTCTTGATTTGGC |
| | Reverse | CAGAAATGATCTTGACGACG |
| Alp5a | Forward | TGTGTCTATGCTTTTCTATCATGC |
| | Reverse | AAGACCGTTGTTTTTATTGCG |

| Primer pair | Direction | Primer sequence (5' - 3') |
|---|---|---|
| 1 | Forward | CTTGATTTGGCATAATACTATGAG |
| | Reverse | GCAGAAATGATCTTGACGACG |
| 2 | Forward | CTTGATTTGGCATAATACTATGAG |
| | Reverse | GAGAGGTGTTAAGCCAGAG |
| 3 | Forward | GGGGTGAGCATTTAAAGC |
| | Reverse | GCAGAAATGATCTTGACGACG |
| 4 | Forward | GTTATGTTTAACAATATACACGC |
| | Reverse | ATATCTCGAGCTATCTTTTAAATGTCATGTAGACA |
| 5 | Forward | CATTCCTTTGTACAGCAACAG |
| | Reverse | TATGTCACAACATCCTACTCC |
| 6 | Forward | CATTCCTTTGTACAGCAACAG |
| | Reverse | GAGAGGTGTTAAGCCAGAG |
| 7 | Forward | GGGGTGAGCATTTAAAGC |
| | Reverse | ATATCTCGAGTTAATACAATAACTTTCCTTGTAAG |
| 8 | Forward | ATATGCTAGCAGCATGAATAAAAAGGACGAAATA |
| | Reverse | ATATCTCGAGTTAATACAATAACTTTCCTTGTAAG |
| 9 | Forward | CTTGATTTGGCATAATACTATGAG |
| | Reverse | ATATCTCGAGCTATCTTTTAAATGTCATGTAGACA |
| 10 | Forward | GTGGAGGTTCCTTGAGTTC |
| | Reverse | GCAGAAATGATCTTGACGACG |
| 11 | Forward | ATATGCTAGCGAAATAAAATCCAAAAAATATGATAAC |
| | Reverse | ATATCTCGAGCTATCTTTTAAATGTCATGTAGACA |
| 12 | Forward | CATTCCTTTGTACAGCAACAG |
| | Reverse | ATATCTCGAGTTAATACAATAACTTTCCTTGTAAG |
| 13 | Forward | CGAAACATGGAGAGAATTAGG |
| | Reverse | TATGTCACAACATCCTACTCC |
